# Whole genome sequencing identifies independent outbreaks of Shigellosis in 2010 and 2011 in La Pampa Province, Argentina

**DOI:** 10.1101/049940

**Authors:** Isabel Chinen, Marcelo Galas, Ezequiel Tuduri, Maria Rosa Viñas, Carolina Carbonari, Anabella Della Gaspera, Daniela Nápoli, David M Aanensen, Silvia Argimón, Nicholas R Thomson, Darren Hughes, Stephen Baker, Caterina Guzmán-Verri, Matthew TG Holden, Alejandra M Abdala, Lucia P Alvarez, Beatriz Alvez, Rosana Barros, Shirley Budall, Constanza Campano, Luciana S Chamosa, Paul Cheddie, Daniel Cisterna, Denise De Belder, Milena Dropa, David Durand, Alan Elena, Gustavo Fontecha, Claudia Huber, Ana Paula Lemos, Luciano Melli, Roxana Elizabeth Paul, Lesly Suarez, Julian Torres Flores, Josefina Campos

## Abstract

*Shigella sonnei* is an emergent cause of diarrheal disease in middle-income countries. The organism causes endemic disease and is also associated with sporadic outbreaks in susceptible populations. In 2010 and 2011 there were two suspected outbreaks of diarrheal disease caused by *S. sonnei* in La Pampa province in central Argentina. Aiming to confirm these as outbreaks and provide insight into the relationship of the strains causing these infections we combined antimicrobial susceptibility testing and pulsed field gel electrophoresis (PFGE) with whole genome sequencing (WGS). Antimicrobial susceptibility testing suggested the two events were unrelated; organisms isolated in 2010 exhibited resistance to trimethoprim sulphate whereas the 2011 *S. sonnei* were non-susceptible against ampicillin, trimethoprim sulphate and cefpodoxime. PFGE profiling confirmed the likelihood of two independent outbreaks, separating the isolates into two main XbaI restriction profiles. We additionally performed WGS on 17 isolates associated with these outbreaks. The resulting phylogeny confirmed the PFGE structure and separated the organisms into two comparatively distantly related clones. Antimicrobial resistant genes were common, and the presence of an OXA-1 was likely associated with resistance to cefpodoxime in the second outbreak. We additionally identified novel horizontally transferred genetic material that may impinge on the pathogenic phenotype of the infecting strains. Our study shows that even with a lack of supporting routine data WGS is an indispensible method for the tracking and surveillance of bacterial pathogens during outbreaks and is becoming a vital tool for the monitoring of antimicrobial resistant strains of *S. sonnei*.

## Report

Dysenteric diarrhea caused by members of the bacterial genus *Shigella* (comprised of the species *S*. flexneri, *S*. sonnei, *S*. boydii and *S*. dysenteriae) remains an on-going public health issue in many industrializing countries. It is estimated that the global burden of disease caused by *Shigella* spp. is ~125 million cases annually [1], the majority of these cases arise in children aged under five years. The Global Enteric Multicenter Study (GEMS), a case-control study of paediatric diarrheal disease conducted in Africa and Asia, found that enterotoxigenic *Escherichia coli* (ETEC) and *Shigella* were the two most common bacterial agents of diarrhea in sub-Saharan Africa and South Asia [2,3]. Notably, *Shigella* spp. were the most prevalent pathogen among children between 24 and 59 months old [3].

Of the four *Shigella* species *S. flexneri* and *S. sonnei* are responsible for the vast majority of the global burden of disease. Traditionally, *S. sonnei* has been the predominant cause of bacterial dysentery in industrialized countries, whereas *S. flexneri* has been considered to be associated with endemic disease and travel to lower income countries [4]. However, this trend is changing as *S. sonnei* is now emerging as a problem in lower middle-income countries, seemingly replacing *S. flexneri* as the leading cause of dysentery in these locations [5]. This trend has also been observed in parts of Latin America [6,7], roughly correlating with improvements in sanitation, water quality and, potentially, a fall in passive immunity against *S. sonnei* via a decline in other bacteria associated with poor water quality [8]

Argentina is a middle-income country in South America with endemic Shigellosis [9], the number of officially reported cases of Shigellosis in 2014 was 4,116. This represents a comparatively high proportion of the 6,200 cases of confirmed the cases of bacterial diarrhea in 2014. *Shigella* outbreaks occur sporadically and rapidly and are frequently associated with changes in antimicrobial susceptibility [10,11]. Aiming to better understand the dynamics of Shigellosis in Argentina we gathered bacterial isolates from two suspected outbreaks of Shigellosis investigated by the public health authorities in Argentina between 2010 and 2011. The two temporally independent events were attributed to *S. sonnei* and occurred within the La Pampa province in the central region of the country. In this retrospective study we combined available epidemiological data and microbiological data with Pulsed Field Gel Electrophoresis (PFGE) and whole genome sequencing (WGS) of the *S. sonnei* isolates from the suspected outbreaks in 2010 and 2011 to investigate their relatedness. We also analysed the discriminatory ability of WGS compared to PFGE, the current international gold-standard method for public health strain tracking.

In December 2009 the Gobernador Centeno hospital in the city of General Pico reported an increased number of cases of diarrhea above the expected endemic rate, an outbreak was suspected. The first *Shigella* (confirmed by standard microbiological methods to be *S. sonnei*) was isolated on the 7^th^ January 2010; the last culture confirmed case of *S. sonnei* was on the 26th February 2010. The cases were distributed throughout (i.e. no apparent case clustering) General Pico. There were 26 reported cases, of which detailed microbiological data was available on nine. Of the 26 reported cases, 13 were in children aged between 0-5 years, and 13 cases were aged between 6-69 years (median: five years); 10 cases were female and 16 were male. No epidemiological association was recorded between cases, apart from a potential cluster in a single household (n=4 cases). The most common disease presentations were diarrhea with blood and mucus (13/26; 50%) and diarrhea with blood without mucus (11/26; 42%). Of the nine available *S. sonnei* organisms isolated during the potential 2010 outbreak, all had an identical antimicrobial susceptibility patterns by disc diffusion [12], exhibiting susceptibility against ampicillin, ciprofloxacin, nitrofurantoin, fosfomycin, naladixic acid and cefpodoxime and non-susceptibility against trimethoprim sulphate.

The second potential outbreak occurred the city of Castex, also in the province of La Pampa (60 Km from General Pico) in the summer of 2011 (3^rd^ February 2011 to 30^th^ March 2011). No supporting epidemiological data were available for this potential outbreak and six *S. sonnei* were isolated. An equal proportion of males and females were infected and the patient age range was 5-26 years (median: eight years). The antimicrobial susceptibility profile of the organisms demonstrated that all organisms were non-susceptible against ampicillin, trimethoprim sulphate and cefpodoxime. Additional laboratory testing suggested that all organisms in this second potential outbreak exhibited AmpC production.

To confirm the likelihood of outbreaks and to investigate the temporal and spatial relationship between organisms we performed PFGE after XbaI digestion on 17 available isolates from 2010 (n=9) and 2011 (n=7) and one additional contextual strain isolated in General Pico in 2013 using standardized PulseNet protocols as previously described (Figure 1) [13]. The XbaI PFGE generated nine differing restriction patterns that could be grouped into two major groups (ARJ16X01.0086 and ARJ16X01.0318) that correlated precisely with their year of isolation; ARJ16X01.0086 is the most frequent pattern described in Argentina. These major restriction patterns differed by six fragments and had 85% pattern similarity. Additional PFGE with BlnI (again following standardized PulseNet protocols [13]) methods of on a limited subsample confirmed this grouping, signifying independent outbreaks likely caused by two differing clones of *S. sonnei* that could be distinguished by their antimicrobial susceptibility patterns. These cases clusters were additionally confirmed using the SatScan function in WHONET software.

**Figure 1.**
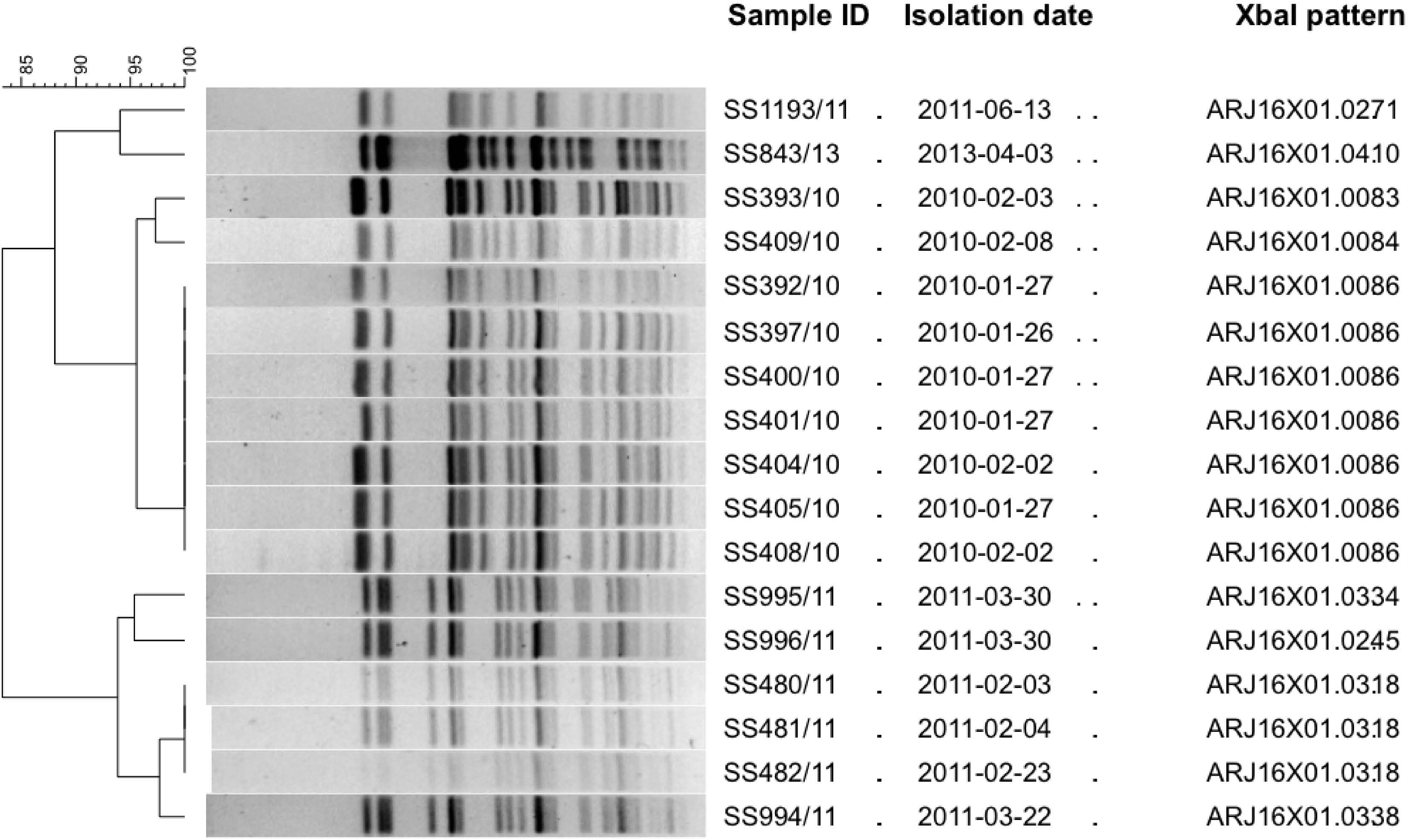
The relationship between *Shigella sonnei* isolated in independent outbreaks in La Pampa province, Argentina in 2010 and 2011. Dendogram based on PFGE profile after XbaI digestion of *Shigella sonnei* isolated from stool samples in 2010 (General Pico) and 2011 (Castex). One additional isolate from 2013 (General Pico) was included as contextual strain. Information regarding the strain ID, the date of isolation and the digestion pattern (according to PulseNet Latinoamerica) are provided.

PFGE does not provide sufficient resolution for phylogenetic inference, to understand the genetic relationship between the organisms from the two outbreaks we performed WGS on nine *S. sonnei* isolated in 2010, seven *S. sonnei* isolated in 2011 and the single contextual isolate from 2013. We identified single nucleotide polymorphisms (SNPs) in comparison to a Chinese reference strain (Ss046, accession number CP000038[14]), as previously described [11,15,16], and constructed a maximum likelihood phylogeny of the 18 strains, identifying approximately 1,500 variable nucleotide sites (Figure 2). Our data confirmed that the outbreaks were associated with two differing, distantly related clones of *S. sonnei*. The first clone (-10 suffix in Figure 2) was comprised of the 2010 isolates; six were highly related, with three remaining isolates in the same group but located on longer branches. The second clone (-11 suffix in Figure 2) contained all six isolates from 2011; these isolates were almost identical, containing less than 10 nucleotide substitutions across their genomes. An additional isolate from 2011 (1193-11) lay outside this group and was deemed not to part of the same clonal outbreak. Notably, the 2011 clone could be distinguished by non-susceptibility against cefpodoxime (yellow nodes in Figure 2) (Data accessible and viewable at http://microreact.org/project/EkJeuWfx-).

**Figure 2.**
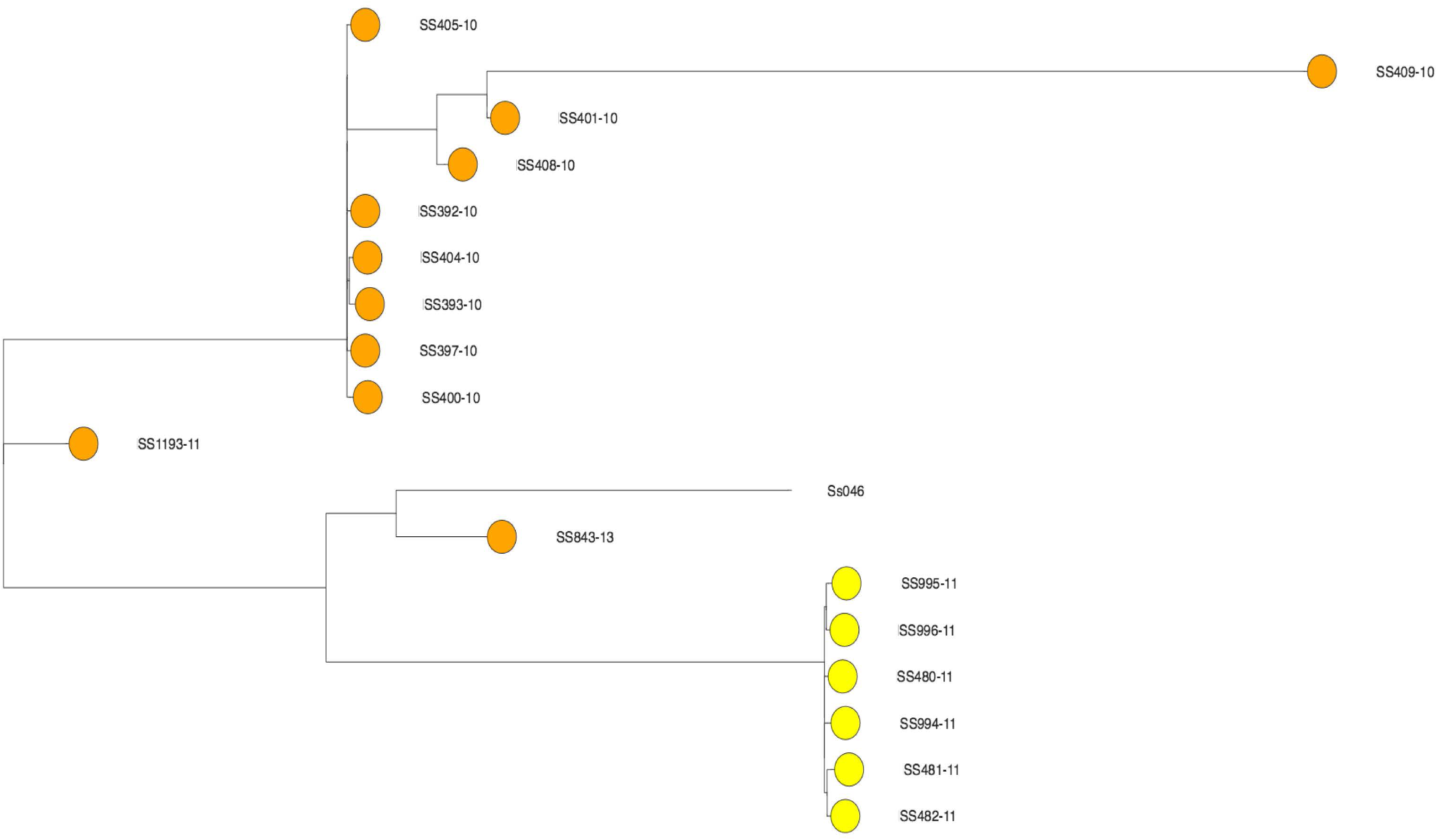
The phylogenetic relationship of *Shigella sonnei* isolated in independent outbreaks in La Pampa province, Argentina in 2010 and 2011. Unrooted maximum likelihood tree constructed using approximately 1,500 variable nucleotide sites from nine organisms isolated in 2010, seven isolates from 2010 and a single isolate from 2013 in La Pampa province, Argentina. *S. sonnei* Ss046 was added as the reference strain. Tree was viewed in microreact and nodes are labelled with the strain name and the year of isolation suffix collared according to susceptibly against cefpodoxime susceptibility (orange; susceptible, yellow, non-susceptible).

We next assembled the genome sequences from the two independent outbreaks to identify additional horizontally transferred genetic material that may be associated with each of the clones and to classify the genes associated with changes in antimicrobial susceptibility. We found that the −10 clone contained a *dfrA1* gene, which is associated with resistance against trimethoprim sulphate. Further, we identified genes associated with resistance to additional antimicrobials that were not tested, including streptomycin (*strAB*), tetracycline (*tetAR*) and sulphonomides (*sulII*). The −11 clone also contained a *dfr* gene (A5) and genes associated with resistance against chloramphenicol (*catA1*), streptomycin (*strAB*), tetracycline (*tetB*) and sulphonomides (*sulII*). We also identified several AmpC β-lactamase genes potentially explaining the non-susceptibility against cefpodoxime including CMY and the Extended Spectrum Beta Lactamase (ESBL) gene, OXA-1. Further, in the −11 clone we identified and assembled a large (>90 Kb) plasmid that exhibited substantial homology and synteny to the recently described 96 Kb p12-4374_96 plasmid in *Salmonella* Heidelberg [17] (accession number: CP012929). This plasmid, not previously described in *Shigella*, encoded a multitude of potentially interesting functions including a conjugation system, a type IVb pilus and the ethanolamine utilization protein, EutE.

Here we have combined traditional methods for tracking bacterial pathogens (antimicrobial susceptibility testing and PFGE) and combined them with WGS to evaluate two potential outbreaks of Shigellosis in a single province in 2010 and 2011 in Argentina. Our data suggests that there were two independent outbreaks of *S. sonnei* induced diarrhea in 2010 and 2011, finding that the organisms causing these case clusters were distantly related to each other. This was somewhat unexpected given the geographical proximity of these two locations and signifies that multiple clones of *S. sonnei* are likely circulating in Argentina, several of which have outbreak potential. We found that the antimicrobial susceptibility profile was sufficient to distinguish between these outbreaks, providing an almost perfect temporal correlation with cefpodoxime resistance. This relationship was further confirmed by PFGE, the current gold standard for strain tracking in such scenarios in Argentina [18]. However, PFGE additionally over predicted the variability within the genomic structures, identifying several banding patterns within the specific clones.

In this particular investigation WGS augmented the findings of the conventional approaches and provided new insight into these outbreaks. Firstly, the phylogenetic inference, which has become standard for WGS of *S. sonnei* [11,15,16], permitted us an exquisite view of the relationship between and within the outbreaks, eventually confirming the two outbreaks. Further, assembly of the genome sequences identified the presence of the range antimicrobial resistance genes, predicting resistance to additional antimicrobials that were not susceptibility tested. These data permitted us to detect the ESBL gene OXA-1 [19], which we hypothesized to be associated with resistance against the third generation cephalosporin, cefpodoxime. ESBL genes are becoming more commonly reported in *Shigella* in Asia [20]. Our data predict that this concerning phenomenon is additionally occurring in Latin America via differing determinants. Oral third generation cephalosporins are one of the current mainstays of treatment for Shigellosis in Argentina, we recommend further genomic surveillance in this region to detect circulating beta lactamase genes. The assembled genome sequences additionally identified further novel sequences encoding potential virulence associated loci that may have a phenotypic effect during infection. These novel genes require conformation and additional experimentation to confirm their role in disease.

Our study contains some limitations including a lack of strain diversity for better phylogenetic inference and a lack of epidemiological data. However, our investigation of these outbreaks represents a “real life” scenario, where limited data hamper contextualization. Here we show that even with a lack of supporting routine data WGS becomes an indispensible method for the tracking and surveillance of bacterial pathogens during outbreaks.

## Acknowledgements

This work was conducted as a component of the genomics and epidemiological surveillance of bacterial pathogens course held from the 17-22 April 2011 in the Dr. Carlos G. Malbran Institute in Buenos Aires, Argentina. We wish to acknowledge the Wellcome Genome Campus Advanced Courses for providing financial and administrative support for the course that supported this investigation, including travel scholarships for attending students. We additionally wish to acknowledge the Dr. Carlos G. Malbran Institute for providing access to these data for training and publication purposes.

